# ProToDeviseR: an automated protein topology scheme generator

**DOI:** 10.1101/2024.06.04.597333

**Authors:** Petar B. Petrov, Valerio Izzi

## Abstract

**Motivation:** Amino acid sequence characterization is a fundamental part of virtually any protein analysis, and creating concise and clear protein topology schemes is of high importance in proteomics studies. Although numerous databases and prediction servers exist, it is challenging to incorporate data from various, and sometimes contending, resources into a publication-ready scheme.

**Results:** Here, we present the Protein Topology Deviser R package (ProToDeviseR) for the automatic generation of protein topology schemes from database accession numbers, raw results from multiple prediction servers, or a manually prepared table of features. The application offers a graphical user interface, implemented in R Shiny, hosting an enhanced version of Pfam’s domains generator for the rendering of visually appealing schemes. Results show that ProToDeviseR can easily and quickly generate topology schemes by interrogating UniProt or NCBI GenPept databases and elegantly combine features from various resources.

**Availability:** Source code (https://github.com/izzilab/protodeviser) is available under GPLv3 license; the tool is also accessible for online use (https://matrinet.shinyapps.io/protodeviser/), without the need of installation.

## Introduction

Proteins are complex molecules, showing an enormous structural and functional versatility (Alberts et al., 2002). Protein topology schemes are a crucial aid to virtually any research in the field of protein analysis and proteomics, as they offer a quick glance into the presence and position of structural domains, regions of functional importance, repeats, motifs, post-translational modifications (PTMs), as well as additional sequence characteristics and peculiarities.

Protein knowledge bases, such as UniProt (The UniProt Consortium, 2023), InterPro (Paysan-Lafosse et al., 2023) and NCBI GenBank (Clark et al., 2016), offer summarized information on numerous protein entries. In addition, various tools for protein features prediction are available, making it possible to complement and extend the characterization of a protein by sequence analysis. Among them are the Simple Modular Architecture Research Tool (SMART) for domain identification (Letunic et al., 2021) and the Eukaryotic Linear Motif (ELM) resource for short functional motif prediction (Kumar et al., 2022). Other resources are dedicated to PTMs, such as NetNGlyc (Gupta and Brunak, 2002) and NetOGlyc (Steentoft et al., 2013) for the detection of N/O-glycosylation, and NetPhos (Blom et al., 2004) and ScanSite (Obenauer et al., 2003) for phosphorylation. Additionally, intrinsically disordered (unstructured) regions of the protein, and segments within them potentially endowed with protein-binding functions, can be investigated with the IUPred/Anchor server (Erdős et al., 2021). Of the three knowledge bases, only UniProt and InterPro offer graphical annotations for proteins and these, though extensive and feature-rich, are poorly suited for direct use in publications due to their spatial organization. On the other hand, feature prediction tools typically focus on simpler visual representations of the results, such as charts, and ignore topology. A notable exception here is SMART, which offers beautiful graphical annotations, albeit restricted almost exclusively to domain organization and lacking, e.g., motifs and PTMs. On the contrary, ELM is able to retrieve information from other resources and plot motif predictions along the protein context, though the verbosity and bundling of the results makes them challenging to incorporate into a publication figure. Finally, a few computational resources for the generation of custom protein schemes exist, such as MyDomains at Prosite (Sigrist et al., 2013) and DOG (Ren et al., 2009), but they require a substantial manual work that significantly hinders their application to projects where many proteins are involved.

Here, we present the Protein Topology Deviser R package (ProToDeviseR) to produce rich, yet concise, protein schemes that are both visually appealing and publication-ready. The application can automatically retrieve information from protein databases, process raw prediction results or a user-provided table of features. ProToDeviseR then automatically transforms the information into a robust annotation code that can be rendered into a topology graph with a single click.

### ProToDeviseR

ProToDeviseR is written in R and its source code is freely available at our GitHub repository under GPLv3 license (https://github.com/izzilab/protodeviser). The application offers a fully functional graphical user interface (GUI), implemented in R Shiny. The function that starts the GUI is called protodeviser_ui and the interface has numerous tooltips and a Help section. In addition to the R package, we also provide a server version, which requires no installation (https://matrinet.shinyapps.io/protodeviser/).

The application generates a code describing protein topology in JSON format, following the syntax used by Pfam’s (Mistry et al., 2021) domain graphics tool. For its annotations, ProToDeviseR searches for specific keywords and classifies protein features into three categories: Regions, Motifs and Markups. The Regions category comprises domains, other relatively long (functional) parts and repeats (Supplementary Fig. S1A). The Motifs category includes short linear motifs, signal peptides, transmembrane parts, as well as disordered regions, compositional biases, coiled-coil regions, low complexity areas, etc (Supplementary Fig. S1B). The Markups category lists single site annotations, such as post-translational modifications or sites of other importance, among which are glycosylations, phosphorylations, active sites, disulfide bonds and many more (Supplementary Fig. S1C). The four internal functions used by the GUI (Fig. 1A), namely id.JSON (generates code from UniProt or NCBI GenPept ID), predicted.JSON (generates code from raw results obtained from various feature prediction resources), custom.JSON (generates code from a user-provided table) and json.TABLE (generates a table from JSON code), can also be used independently and incorporated into other scripts, for example for batch analysis.

**Fig. 1.**
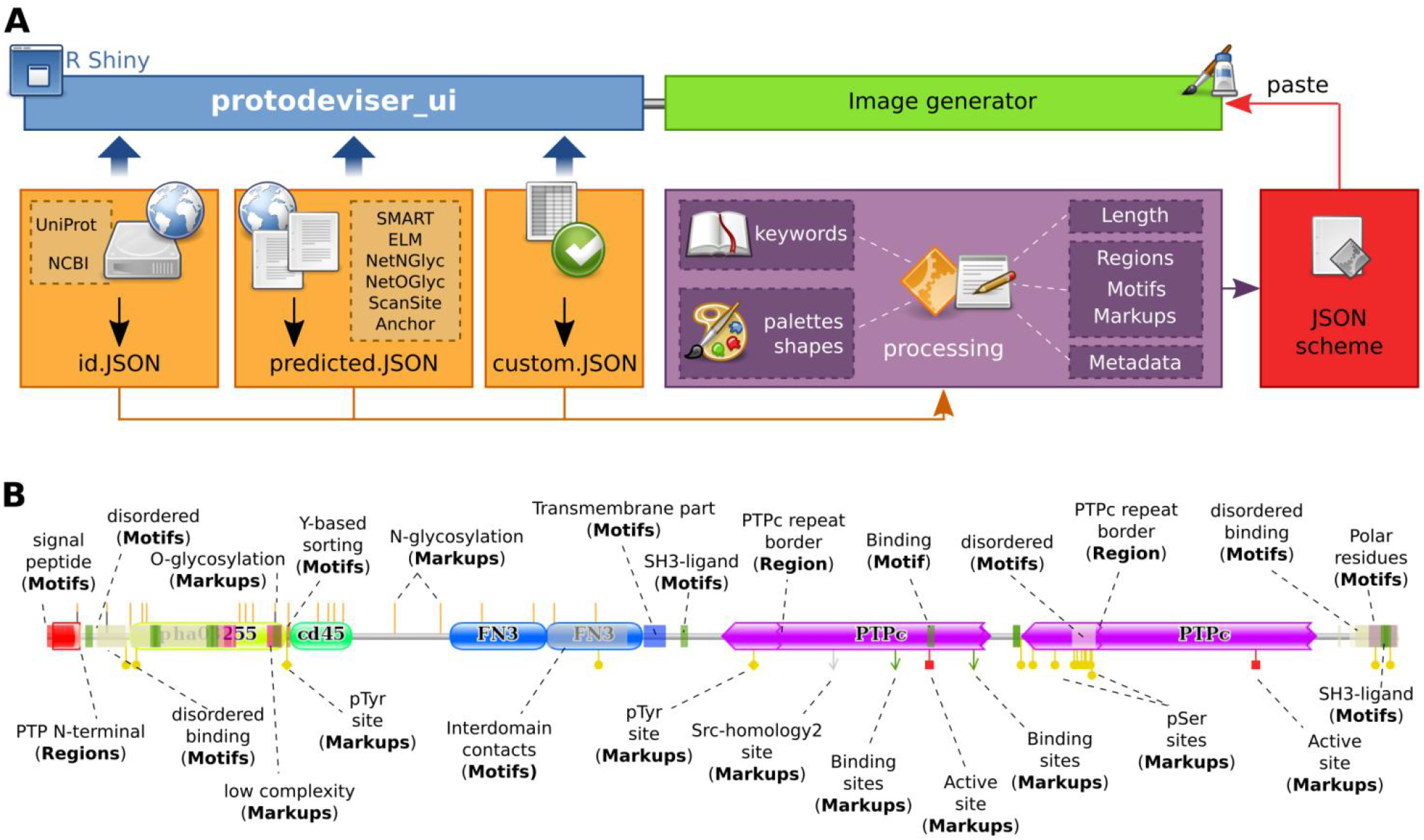
ProToDeviseR generation of protein scheme. **A) Application overview.** The graphical user interface protodeviser_ui (blue), runs on top of the core functions id.JSON, predicted.JSON, custom.JSON (orange), which process (purple) the input to produce a scheme in JSON code (red). Code can be pasted in the Image generator tab of the GUI (green). **B) Example scheme**. CD45 features were extracted from UniProt, NCBI GenPept, as well as predictions were done at SMART, ELM, NetNGlyc, NetOGlyc, NetPhos, ScanSite and IUPred/Anchor (See Supplementary Figure S2). Results were compared and a composite table was prepared and submitted to ProToDeviseR. Protein features are pointed and their respective categories are indicated in brackets.

The online and local implementation of the GUI offer identical views, with an input panel divided into two sections, “Protein ID” and “Protein features”, the latter subdivided into “Predicted” and “Predefined” (Supplementary Fig S2A-D). The “Protein ID” section offers streamlined access to ProToDeviseR functionalities, as it simply requires a UniProt or NCBI GenPept identifier, after which all the necessary data will be automatically imported and prepared. The “Protein features” section offers a more granular approach to input protein data, with the “Predicted” tab accepting predictions from SMART, ELM, NetNGlyc, NetOGlyc, NetPhos, ScanSite and IUPred/Anchor, each settable with own cut off values, as well as fields for protein length and optional metadata, such as protein name. Alternatively, the “Predefined” tab accepts a user-curated table of protein features as typical in data mining, when a list of manually curated features is to be visualized. Upon successful integration of the input(s), the following outputs become available:

∘ Table preview: results are shown as a table which can be inspected, sorted by column and downloaded.
∘ JSON output: the generated topology is shown in a code box, with a download option should further manual changes be desired. In general, users will only need to press the button to copy the contents of the box to the clipboard, so that it can be pasted into the neighbouring tab, called Image generator.
∘ Image generator: pasting the JSON code here will render the graphics. It is important to notice that this part of the application is not directly developed by us, as it is a port of the legacy custom domains generator from Pfam. We have embedded it into ProToDeviseR for user convenience along with a few enhancements, such as proportional image size rescaling, tunable amino acid pixel size and motif opacity. In particular, adjusting the amino acid size zooms the scheme (actually, making it longer), improving its resolution, a particularly useful “trick” for short or feature-rich proteins.

As an example, we devised topological schemes for human CD45 (Receptor-type tyrosine-protein phosphatase C), inputting either the UniProt ID P08575 or the NCBI GenPept ID NP_002829.3 into the “Protein ID” tab, as the general user would (Supplementary Fig. S3A and B). To test for feature integration, we submitted the amino acid sequence of CD45 (1306 aa) to SMART, ELM, NetNGlyc, NetOGlyc, NetPhos, ScanSite and IUPred/Anchor, and fed the results to the “Protein features / Predicted” tab of ProToDeviseR (Supplementary Fig. S2C). Finally, to test the curated table entry functionality, we merged the results from the previous runs, removed redundant entries and submitted the combined table (Supplementary Table S3) to the “Protein features / Predefined” tab of ProToDeviseR (Fig. 1B). Our example data combined the information already available at UniProt and NCBI and added putative novel features in addition to the ones listed at the databases. All the examples files are available from the Help section of the GUI.

## Conclusion

ProToDeviseR offers a fast and easy-to-use interface to comprehensively annotate proteins and devise topological schemes. It seamlessly integrates input from resources that provide only a limited visual representation of their data, or none at all. As a result, ProToDeviseR produces aesthetically pleasant, publication-ready graphs.

### Development

ProToDeviseR was developed in R v4.4.0 (https://www.r-project.org/) running on CRUX v3.7 (https://crux.nu/) distribution of GNU/Linux. All necessary software was installed from the ports freely available at the distribution’s port database (https://crux.nu/portdb/). Figures were assembled in Inkscape (https://inkscape.org/) and icons are from the Tango Desktop Project v0.8.90.

## Acknowledgements

We thank Anjani Fowdar from University of Cape Town (RSA), for preliminary testing of ProToDeviseR.

## Supplementary information

Supplementary data for ProToDeviseR v1.0. For extended, and potentially updated information, consult with the Help section tab of the GUI (protodeviser_ui).

## Supplementary figures

**Fig. S1.**
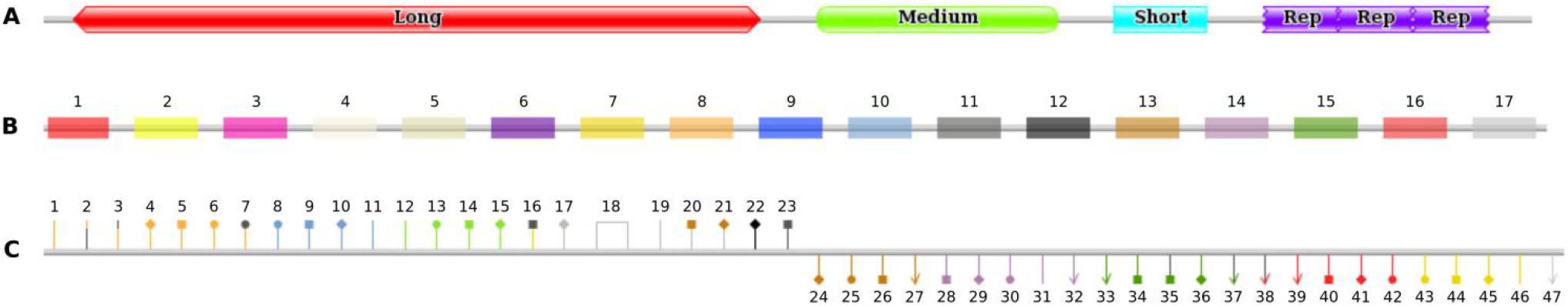
Protein features annotations. **A) Regions**. Long, medium, short and three repeats are shown. At the moment, ProToDeviseR assigns regions colours anew for each protein. A rainbow gradient is used, where regions with the same name will have the same colour. Please note that, although coloured the same within one protein, regions may have another colour assigned to them in another protein. Shapes are: pointed: long regions (> 200 amino acids), curved: medium-sized regions (35-200 amino acids), straight: small regions (< 35 amino acids), jagged: regions designated as repeats. **B) Motifs**. Signal peptide (1), Coiled coil (2), Low complexity (3), Intrinsic disorder (4), Intrinsically disordered binding (5), Charged or polar amino acids patch (6), Phosphorylation motif (7), Glycosylation motif (8), Transmembrane part (9), Lipidation motif (10), Cleavage motif (11), Degradation motif (12), Targeting motif (13), Nuclear localisation or export motif (14), Docking, ligand or binding motif (15), Activity-related motif (16), Other motif (17). **C) Markups**. N-glycosylation (1), O-glycosylation (2), Glycosaminoglycan (3), C-mannosylation (4), O-fucosylation (5), Glycosylation unspecified (6), Hydroxylation (7), Prenylated (8), Acylated (9), GPI (10), Lipidation (11), Acetylation (12), Methylation (13), Amidation (14), Pyrrolidone carboxylic acid (15), Sulfation (16), D-isomerization (17), di-Sulfide bond (18), Cross-linking (19), Sumoylation (20), Ubiquitination (21), Degradation (22), Cleavage (23), Sorting (24), Targeting (25), Retaining (26), Absorption (27), Nuclear import (28), Nuclear export (29), Nuclear receptor (30), Nuclear-related (31), DNA-binding (32), Binding site (33), Ligand binding (34), Ligand site (35), Docking (36), Interacts with (37), Flavin-binding (38), Co-factor (39), Active site (40), Catalytic activity (41), Activity regulation (42), Phospho-Serine (43), Phospho-Threonine (44), Phospho-Tyrosine (45), Phosphorylation unspecified (46), Other motif (47).

**Fig. S2.**
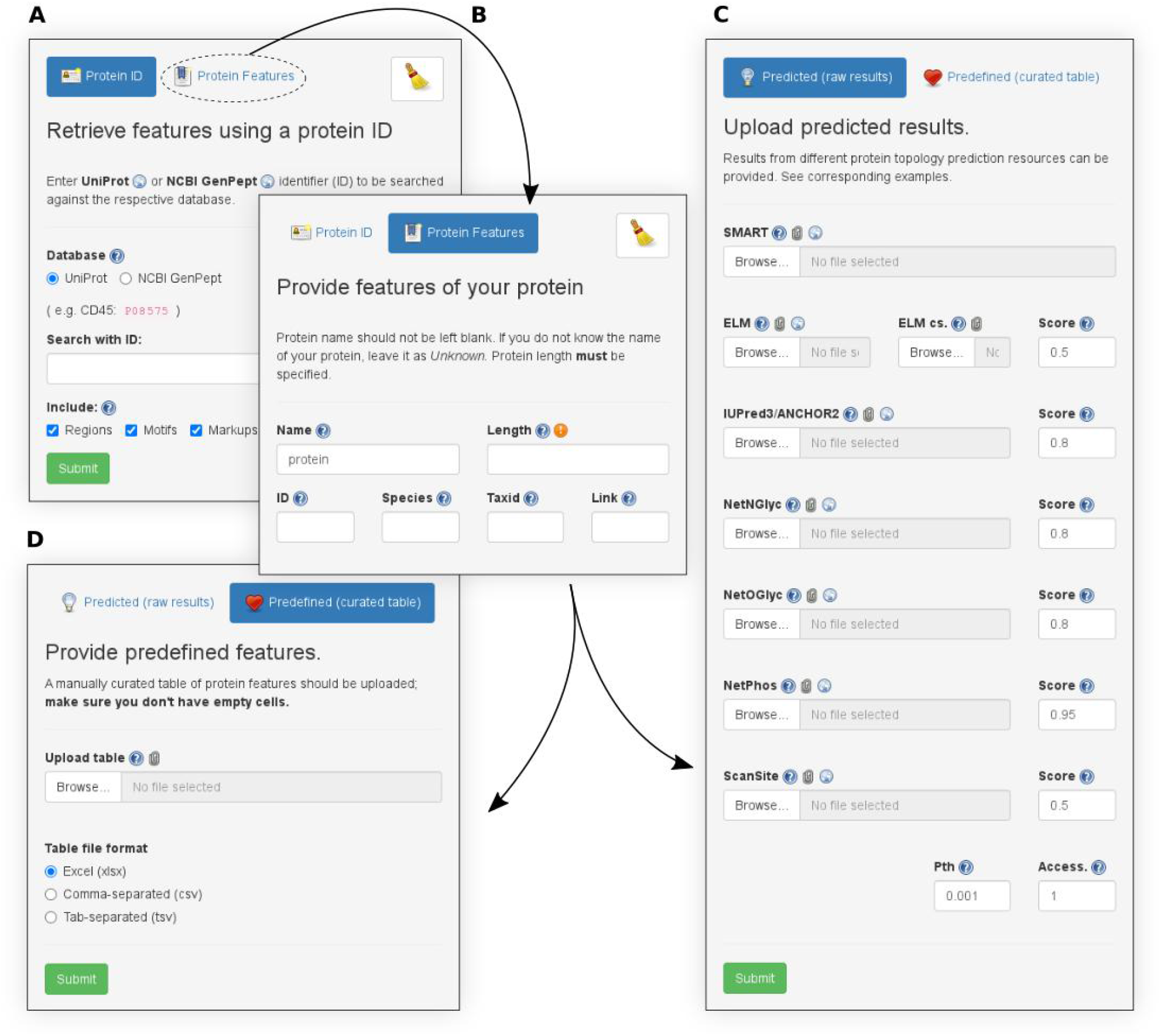
Input panel options. **A) Protein ID**. Simply enter a UniProt or NCBI GenPept ID to be searched. By default, the program will search for regions, motifs and markups, but these can be deselected if desired. **B) Protein Features**. Enter basic metadata information about the protein -- size in aa (required), name, etc. This is used for either C or D. **C) Protein Features / Predefined (raw results)**. The results from several prediction resources can be uploaded here. Threshold cut off values can be specified, where applicable. **D) Protein Features / Predefined (curated table)**. Provide a table of user-defined protein features. Accepted formats are xlsx, csv and tsv.

**Fig. S3.**
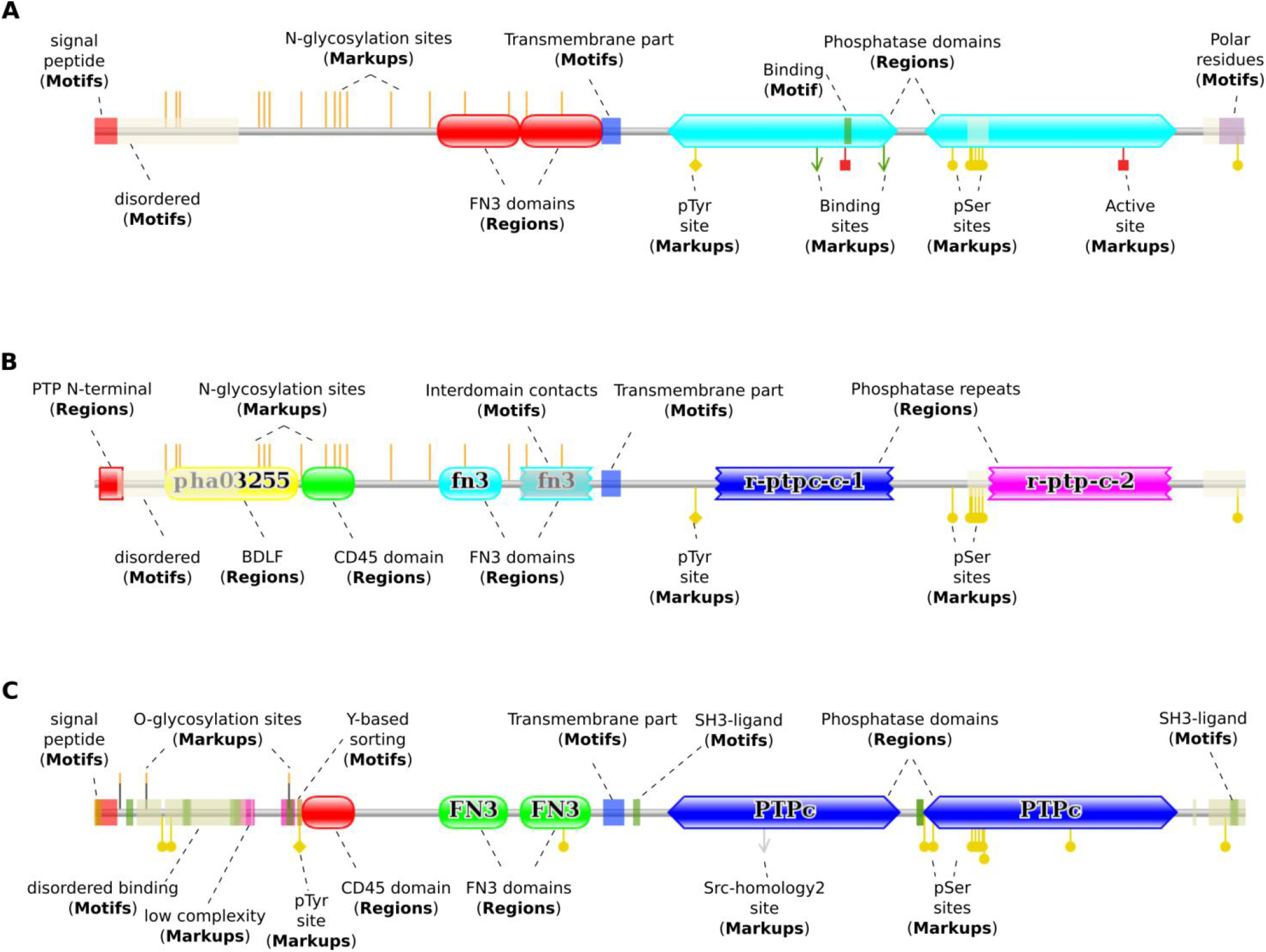
Schemes of CD45 produced by different search strategies of ProToDeviseR. A) The “Protein ID” tab was used with ID “P08575” to retrieve information about the protein from UniProt. B) The “Protein ID” tab was used with ID “NP_002829.3”, analogously for NCBI GenPept. The database denotes the second FN3 domain and the phosphatase domains as repeats, hence the jagged edges. C) Predicted features for sequence with Uniprot ID “P08575”, were submitted at the “Protein features / Predicted” tab. Cut off values were as follows: SMART (NA), ELM (Score > 0, no filtering applied in order to show all motifs), NetNglyc (Score > 0.95), NetOGlyc (Score > 0.95), NetPhos (Score > 0.99), ScanSite (Score > 0.5, Percentile threshold > 0.001, Accessibility > 1) and IUPred3/Anchor2 (Score > 0.6).

## Supplementary tables

**Table S1.** Columns for a user-prepared table with protein topology annotations.

| Motif class | Keywords |
| --- | --- |
| activity_regulation | co-factor, active site, catalytic activity, activity regulation, activity |
| charged_polar_reg | basic, acidic, charged, polar |
| cleavage | cleave, cleavage |
| coiled_coil | coiled coil |
| degradation | degrad, degran, destruct |
| disordered_region | disorder |
| disor_region_bind | disorder bind |
| docking_ligand | ligand, bind, docking, interact |
| glycosylation | glycosaminoglycan, mucopolysaccharide, O-fucos, C-mannos, N-glyc, O-glyc, glycosylation |
| lipidation | prenyl, isopren, farnes, geranyl, dolichol, caax, acylat, myrist, palmit, gpi, glycosylphosphatidylinositol, phosphoethanolamine, lipid |
| low_complexity | low complex |
| nuclear_related | nucleus, nuclear import/localization, nuclear export/localization, zink finger, dna |
| phosphorylation | ser, thr, tyr, phospho, kinase |
| signal_peptide | signal, signal peptide |
| targeting | sumo, absorb, absorption sort, target |
| transmembrane | trans-membrane |

**Table S2.** Keywords used to classify markups.

| Markup class | Keywords |
| --- | --- |
| absorption | absorption, absorb |
| acetylation | acetyl |
| active_site | activ site |
| activity_regulation | activ regulat |
| acylated | acylat, myrist, palmit |
| amidation | amide, amidation |
| binding_site | bind |
| catalytic_activity | catal activ |
| cleavage | cleave, cleavage |
| C_mannosylation | C-mannos |
| cofactor | co-factor |
| cross_link | cross-link |
| degradation | degrad, destruct, degran |
| diSulfide_bridge | disulf |
| dna_binding | dna bind |
| docking | docking |
| flavin_binding | flavin, flavo, fmn, fad |
| glycosaminoglycan | glycosaminoglycan |
| glycosylation | glycosylation, mucopolysaccharide |
| gpi | gpi, glycosylphosphatidylinositol, phosphoethanolamine |
| hydroxylation | hydroxylation, hydrocxy proline/lysine/phenylalanine/arginine/asparagine/aspartate |
| interacts_with | interact |
| isomerization | isomerization, D- |
| ligand_binding | ligand bind |
| ligand_site | ligand |
| lipidation | lipid |
| methylation | methyl |
| N_glycosylation | N-glyc |
| nuclear_export | nucleus/nuclear export |
| nuclear_import | nucleus/nuclear import/localization |
| nuclear_receptor | nucleus/nuclear receptor |
| nuclear_related | nucleus, nuclear |
| O_fucosylation | O-fucos |
| O_glycosylation | O-glyc, O-GalNAc, O-mucin |
| phosphorylation | phosphorylation |
| PhosphoSerine | Ser-phospho, Ser-kinase |
| PhosphoThreonine | Thr-phospho, Thr-kinase |
| PhosphoTyrosine | Tyr-phospho, Tyr-kinase |
| prenylated | prenyl, isopren, farnes, geranyl, dolichol, caax |
| pyrrolidone | pyrrolidone, pyroglutamic, pyroglutamate |
| retaining | retain |
| sorting | sort |
| sulfation | sulfation, sulphotyr, sulphothr, sulphoser |
| sumo | sumo |
| targeting | target |
| ubiquitin | ubiquitin |

**Table S3.** Columns for a user-prepared table with protein topology annotations.

| Column name | Description |
| --- | --- |
| type | classification of the feature (string). Accepted are: regions, motifs or markups. |
| start | start coordinate (numeric) |
| end | stop/end coordinate (numeric). If you denote a markup site, use the same coordinate for start and end. |
| text | short name of feature (string). |
| description | longer description of feature (string). |
| scoreName | if feature was predicted, name of the prediction score (string). Optional |
| score | actual score value (numeric). Optional |
| database | source of feature information (string). Optional |
| accession | database identifier of feature (string). Optional |
| sequence | actual sequence of feature, as amino acids or regex (string). Optional |
| target | if a feature interacts with another partner, indicate partner (string). Optional |

